# Estimating Phylogeny from microRNA Data: A Critical Appraisal

**DOI:** 10.1101/003921

**Authors:** Robert C. Thomson, David C. Plachetzki, D. Luke Mahler, Brian R. Moore

**Affiliations:** Department of Biology, University of Hawai‘i at Mānoa, Honolulu, HI 96822, USA; Department of Molecular, Cellular, and Biomedical Sciences, University of New Hampshire, Durham, NH 03824, USA; Department of Evolution and Ecology, University of California, Davis CA 95616 USA

**Keywords:** miRNA, homoplasy, secondary loss, stochastic Dollo

## Abstract

As progress toward a highly resolved tree of life continues to expose nodes that resist resolution, interest in new sources of phylogenetic information that are informative for these most difficult relationships continues to increase. One such potential source of information, the presence and absence of microRNA families, has been vigorously promoted as an ideal phylogenetic marker and has been recently deployed to resolve several long-standing phylogenetic questions. Understanding the utility of such markers for phylogenetic inference hinges on developing a better understanding for how such markers behave under suitable evolutionary models, as well as how they perform in real inference scenarios. However, as yet, no study has rigorously characterized the statistical behavior or utility of these markers. Here we examine the behavior and performance of microRNA presence/absence data under a variety of evolutionary models and reexamine datasets from several previous studies. We find that highly heterogeneous rates of microRNA gain and loss, pervasive secondary loss, and sampling error collectively render microRNA-based inference of phylogeny difficult, and fundamentally alter the conclusions for four of the five studies that we re-examine. Our results indicate that miRNA data have far less phylogenetic utility in resolving the tree of life than is currently recognized and we urge ample caution in their interpretation.

---

As genomic tools and affordable DNA sequencing have become widely available, our ability to leverage molecular sequence data to estimate species phylogeny has rapidly increased. The flood of molecular data has, in turn, witnessed brisk progress in resolving the tree of life (Sanderson 2008; Thomson and Shaffer 2010). Nevertheless, many relationships have resisted resolution despite repeated efforts using increasing amounts of sequence data. These challenging cases have motivated the search for new sources of (molecular) phylogenetic information, which places precedence on data that evolve by rare and nearly irreversible genomic changes. Patterns of gene rearrangement, duplication, insertion and deletion, as well as positional information for retrotransposons have all been promoted as candidate data with “ideal” phylogenetic properties (*e.g.*, Hillis 1999; Rokas and Holland 2000; Boore 2006; Boore and Fuerstenberg 2008). Although new types of phylogenetic data may hold promise in resolving difficult nodes in the tree of life, they require careful consideration in order to appropriately model the underlying evolutionary process by which they arose and to accommodate possible sampling biases associated with their collection.

One recently promoted class of putatively ideal phylogenetic data is the presence/absence of microRNA (miRNA) families (Dolgin 2012; Tarver et al. 2013). MicroRNAs are small regulatory RNA molecules that play a pervasive role in gene regulation and are understood to influence a variety of biological processes both in normal physiological and pathological disease contexts (Lu et al. 2005; Alvarez-Garcia and Miska 2005). Because of their widespread importance in regulating gene networks and their potential role in the evolution of complexity, miRNAs are currently the subject of considerable focus in developmental biology (Berezikov 2011; Peterson et al. 2009; Heimberg et al. 2008).

The justification for the phylogenetic utility of miRNA presence/absence data stems from the way that novel miRNA families arise. MicroRNAs are believed to originate largely from random hairpin sequences in intronic or intergenic regions (typically 60–80 bp in length) of the genome that become transcribed into RNA (Nozawaet et al. 2010; Campo-Paysaa et al. 2011). After transcription, the resulting primary miRNAs may fold into hairpins that serve as the substrate for a pair of enzymes—called *Drosha* and *Dicer*—involved in miRNA synthesis (Krol et al. 2010), culminating in a mature miRNA (typically 22 bp in length).

The odds that any individual hairpin structure will acquire the requisite mutations to form a novel miRNA are exceedingly slim; however, genomes contain many thousands of these structures, such that novel miRNAs are likely to accumulate over deep time (Nozawaet et al. 2010). After the introduction of new functional miRNAs, strong purifying selection associated with their regulatory role can lead to both extraordinarily low rates of substitution within miRNA sequences, as well as long-term preservation of miRNAs in the genome (Nozawaet et al. 2010). This biological scenario is expected to lead to an evolutionary pattern wherein new miRNAs—over long time scales— continually arise in genomes and experience a low rate of secondary loss (Campo-Paysaa et al. 2011). Moreover, the origin of novel miRNAs involves the accumulation of random mutations to a relatively long sequence (60–80 bp in animals), rendering it highly improbable that identical miRNAs will evolve convergently (Sperling and Peterson 2009). These considerations have led to the promotion of miRNAs as a new source of data that are ideal for parsimony inference of phylogeny: they should exhibit extraordinarily low levels of homoplasy (*i.e.*, they are not expected to arise convergently or to be lost secondarily) and thus provide unambiguous synapomorphies (shared-derived character states) that elevate miRNAs to “one of the most useful classes of characters in phylogenetics” (Heimberg et al. 2010).

The above reasoning has led to a recent proliferation of miRNA-based phylogenetic studies seeking to unequivocally resolve several recalcitrant relationships in the tree of life. At the time of our analysis, these include five formal^1^ phylogenetic analyses of miRNA data focused on identifying the phylogenetic position of turtles within amniotes (Lyson et al. 2011), acoelomorph flatworms within animals (Philippe et al. 2011), lampreys within vertebrates (hagfish and jawed vertebrates; Heimberg et al. 2010), myzostomidan worms within bilaterians (Helm et al. 2012), and to establish the monophyly of—and resolve relationships within—annelids (Sperling et al. 2009).

These studies proceed by first identifying the set of miRNAs present in each study lineage using one of two general approaches: by searching for known or novel miRNAs either in existing genome assemblies and/or in novel data generated by sequencing small-RNA libraries. The identified miRNA families are then used to construct a data matrix in which each miRNA family is treated as an ordered binary character, where miRNA presence is the derived state. Finally, this data matrix is subjected to (Dollo or Wagner) parsimony analysis to estimate phylogenetic relationships.

Here, we critically examine the use of miRNA data for phylogeny estimation, focusing on three concerns: 1) the validity of claims related to the evolution of miRNA families (*i.e.*, that secondary loss is exceptionally rare); 2) limitations of parsimony methods used to infer phylogeny from miRNA presence/absence data; and 3) problems associated with the detection of miRNA families. We demonstrate that these concerns collectively render published phylogenetic conclusions based on miRNA data uncertain (obscured by their reliance on non-statistical methods) and/or strongly biased (owing to miRNA-detection problems and/or inference method). We illustrate these concerns by reanalyzing five published phylogenetic studies of miRNA data.

## Interpreting and analyzing microRNA data: Is miRNA absence evidence or absence of evidence?

In order to properly analyze and interpret miRNA presence/absence data, we must be explicit on the nature and meaning of *absence*. A microRNA family that is scored as absent in a particular lineage can, in principle, have one of three histories: 1) the miRNA family may have never arisen in or been inherited by that lineage (‘true absence’); 2) the miRNA family may have previously been present in the lineage but subsequently lost from the genome (‘secondary loss’); or 3) the miRNA family may actually be present in the genome but escaped detection during data collection (‘sampling error’). If all (or nearly all) absences of miRNA families are true absences, then miRNA loss strictly does not occur (or occurs exceedingly rarely): this is the (implicit) assumption of miRNA studies. Accordingly, because the evolution of miRNA data involves minimal character change—miRNA families have a unique origin (bereft of convergence) with negligible/no secondary loss—the use of parsimony as an inference method might be justified.

In fact, nearly all published miRNA studies (including all five re-examined here) have used some variant of the parsimony method to estimate phylogeny. The miRNA study by (Sperling et al. 2009) used “standard” (Wagner) parsimony—in which gains and losses of miRNA families incur equal cost (Kluge and Farris 1969), and the remaining four studies (Heimberg et al. 2010; Lyson et al. 2011; Philippe et al. 2011; Helm et al. 2012) employed Dollo parsimony (LeQuesne 1974). Dollo parsimony allows for the unique evolution of a character and its subsequent loss (both with equal cost), but precludes re-evolution of the same character (with effectively infinite cost) once it has been lost.

## Secondary loss of miRNA families is (apparently) common

Here we explore the claim that secondary loss of miRNA families is exceedingly rare (*e.g.*, Sempere et al. 2007; Sperling and Peterson 2009; Wheeler et al. 2009). To this end, we derived estimates of the prevalence of miRNA loss from analyses of published miRNA datasets. The prediction is quite simple: if loss of miRNA families is exceedingly rare, then the most parsimonious tree for a given miRNA dataset should be (virtually) free of homoplasy (implied secondary loss of miRNA families), given that Dollo parsimony does not permit convergent or parallel evolution.

To derive estimates of the implied prevalence of miRNA loss, we reanalyzed the miRNA datasets under Dollo parsimony with PAUP* v4b10 (Swofford 1998) by means of exhaustive searches, treating all characters as ‘Dollo.up’, which provides the parsimony score (*i.e.*, the total number of implied miRNA gains and losses) for the optimal tree. We then tabulated the number of miRNA losses using the ‘dollop’ function in Phylip v3.5c (Felsenstein 1993). Finally, we estimated the prevalence of miRNA secondary loss in each of the five formal miRNA phylogenetic studies, which is simply calculated as the number of implied losses divided by the parsimony score (total number of implied changes).

Our survey of published studies suggests that secondary loss of miRNA families is apparently quite common (Table 1). In all but one study (Lyson et al. 2011; addressed below), secondary miRNA losses constitute between 27–54% (with an overall average of 38%) of the implied evolutionary changes. These phylogenetic results accord well with those of molecular evolutionary studies, in which prevalent secondary loss of miRNA families have been inferred for various taxa (Nozawaet et al. 2010; Guerra-Assunçáo and Enright 2012; Meunier et al. 2013; Lyu et al. 2014).

**Table 1:** Prevalence of miRNA loss inferred under Dollo parsimony and the stochastic Dollo model.

| miRNA study | Reference | # Parsimony informative | Optimal parsimony score<sup>a</sup> | # implied miRNA losses<sup>b</sup> | Proportion of secondary loss<sup>c</sup> | Estimated rate of miRNA loss (mean, [HPD])<sup>d</sup> |
| --- | --- | --- | --- | --- | --- | --- |
| Amniotes<sup>e</sup> | Lyson et al. (2011) | 34 | 36 | 1 | 0.03 | $1.99 \times 10^{-4}$, [$3.48 \times 10^{-6}$, $4.75 \times 10^{-4}$] |
| Animals | Philippe et al. (2011) | 115 | 158 | 43 | 0.27 | $2.01 \times 10^{-4}$, [$4.05 \times 10^{-6}$, $4.79 \times 10^{-4}$] |
| Annelids<sup>f</sup> | Sperling et al. (2009) | 71 | 113 | 42 | 0.37 | $1.99 \times 10^{-4}$, [$9.15 \times 10^{-6}$, $4.75 \times 10^{-4}$] |
| Bilaterians | Helm et al. (2012) | 71 | 147 | 79 | 0.54 | $2.01 \times 10^{-4}$, [$2.73 \times 10^{-6}$, $4.82 \times 10^{-4}$] |
| Vertebrates | Heimberg et al. (2010) | 172 | 249 | 84 | 0.34 | $2.04 \times 10^{-4}$, [$1.08 \times 10^{-5}$, $4.87 \times 10^{-4}$] |

<sup>a</sup>Parsimony optimal trees were inferred with PAUP\*v4b10 (Swofford, 2003) by means of exhaustive searches, treating all characters as ‘Dollo.up’, which logs the number of parsimony-informative characters and the parsimony score.
<sup>b</sup>The number of implied miRNA losses was tabulated using the ‘dollop’ function in Phylip v3.5c (Felsenstein, 1993).
<sup>c</sup>The number of implied losses divided by the parsimony score (total number of implied miRNA gains and losses).
<sup>d</sup>The mean and [lower, upper] 95% highest posterior density (HPD) estimates of the marginal posterior probability density for the rate of miRNA loss were inferred using BEAST v1.7.5 (Drummond et al. 2012) under the stochastic Dollo model (Nicholls and Gray, 2008; Alekseyenko et al., 2008) and the optimal relaxed-clock model without topological constraints.
<sup>e</sup>The number of implied miRNA losses calculated here (and reported in the original study) is an underestimate. The original study indicates that additional miRNAs were detected that entailed secondary losses (see supplementary Table 1, Lyson et al., 2011), but these data were excluded from the dataset.
<sup>f</sup>The original study (Sperling et al., 2009) for this dataset used ‘standard’ (Wagner) parsimony, which implies a greater degree of secondary miRNA loss (results not shown).

Although we suspect that the degree of secondary loss in published studies is somewhat inflated by miRNA sampling errors (see: *Sampling error in miRNA detection and its phylogenetic impact*, below), the complex character histories of miRNA evolution nevertheless suggest that the use of parsimony—which effectively places all of the probability on the single character history with the absolute minimal amount of change—is not a suitable method with which to infer phylogeny from miRNAs.

## Statistical analysis of miRNA exposes considerable phylogenetic uncertainty

As discussed in the preceding section, the evolution of miRNA often appears to be complex, which raises concerns about the choice of parsimony as a method of inference. Stochastic models are available that are more appropriate for accommodating complex histories, as the likelihood of a given character (in this case, a miRNA family) is calculated by integrating over all possible character histories (in this case, patterns of miRNA gain and secondary loss that could give rise to the observations), weighting each history by its probability under the model. Furthermore, stochastic models are available that may be appropriate for the analysis of miRNA presence/absence data. For example, the binary stochastic Dollo model (SD: Nicholls and Gray 2008; Alekseyenko et al. 2008) appears to be well suited for the analysis of miRNA presence/absence data. The SD model describes an immigration-death stochastic process in which the origin of a character (miRNA family) is modeled as a homogeneous Poisson process with instantaneous rate *λ*, and its subsequent loss is modeled as a stochastic branching process (where the probability of loss is proportional to the branch length in which it persists toward the present) with an instantaneous rate of secondary loss, *µ* (Alekseyenko et al. 2008). Inference under stochastic models within a Bayesian statistical framework provides a natural means for assessing support/accommodating uncertainty in phylogenetic estimates. Because the majority of published miRNA studies to date have either ignored the issue of evidential support for estimates, or have relied on *ad hoc* support measures (such as the Bremer support index; Bremer 1988) which have no clear statistical interpretation, the availability of an inference framework that explicitly assesses support is particularly attractive.

Markov chain Monte Carlo (MCMC) simulation is used to approximate the joint posterior probability distribution of the phylogenetic parameters. A Markov chain is specified that has state space comprising all possible values for the phylogenetic model parameters, which has a stationary distribution that is the distribution of interest (*i.e.*, the joint posterior probability distribution of the model parameters). Samples drawn from the stationary Markov chain provide valid estimates of the joint posterior probability density, which can be queried marginally with respect to any parameter of interest. In the case of topology, the marginal posterior probability for a given clade is simply its frequency in the sampled trees.

*Bayesian inference of phylogeny from miRNA datasets.*—These considerations motivated us to re-analyze previously published miRNA datasets within a Bayesian statistical framework using a stochastic binary Dollo model (Alekseyenko et al. 2008) to describe the gain and loss of miRNA families. For each of the five miRNA datasets, we treated all characters as ‘Dollo type’ and approximated the joint posterior probability density via MCMC using BEAST v1.7.5 (Drummond et al. 2012). We specified a prior for the rate of miRNA loss, *µ*, using an exponential distribution with a small rate parameter 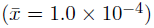 and specified a prior on the tree topology and node heights using a stochastic birth-death branching process.

Molecular studies have alternatively characterized the evolution of miRNAs as a gradual process of continuous accumulation via mutation (Nozawaet et al. 2010), or as an episodic process associated with major regulatory or developmental innovations (Campo-Paysaa et al. 2011). Accordingly, we explored an array of (relaxed) clock models to describe the variation in rates of miRNA evolution across the tree or through time that range from stochastically constant to episodic. Specifically, for each dataset, we performed analyses under the strict-clock model, the random-local clock model (RLMK: Drummond and Suchard 2010), and the uncorrelated lognormal (UCLN) and exponential (UCED) relaxed-clock models (Drummond et al. 2006). Inference of the joint posterior probability density for each composite phylogenetic model (*i.e.*, the binary stochastic Dollo model + one of the [relaxed] clock models) involved at least three independent MCMC analyses, running each chain for 100 million cycles and sampling every 10,000*^th^* cycle.

In order to compare fit of the data to these four alternative clock models, we performed additional analyses targeting the marginal likelihood of the data under each of the four composite phylogenetic models. For each dataset, this entailed running the MCMC through a series of 50 power posteriors spanning from the prior to the posterior, with the powers spaced along a Beta(0.3, 1.0) distribution. We then estimated the marginal likelihood from this chain using both path and stepping stone sampling analyses (Baele et al. 2012). These analyses were also each repeated at least three times to ensure stability of the marginal likelihood estimates. We then compared support for the alternative clock models by calculating Bayes factors as the ratio of the marginal likelihoods for each pairwise combination of candidate models. We interpret Bayes factors following Kass and Raftery (1995): viewing 2 ln BF values >10 as very strong support for the candidate model, between 6 and 10 as strong support, between 2 and 6 as positive evidence, and < 2 as essentially equivocal regarding the alternative models. We performed model comparison only for models where the analyses performed very well, judged by the MCMC mixing efficiently across the power posteriors and highly stable estimates of the marginal likelihood across replicated analyses with both stepping stone and path sampling.

In total, this analysis design entailed 180 MCMC analyses: each of the five miRNA datasets were analyzed under each of the four (relaxed) clock models, performing three independent MCMC analyses under each model, repeating analyses to target first the joint prior probability, then the joint posterior probability, and finally the marginal likelihood densities. We assessed the performance of each MCMC analysis for all parameters (including the topology) using Tracer and AWTY (Rambaut and Drummond 2007; Nylander et al. 2008), which suggested that the chains mixed well and had converged prior to *∼* 50 million cycles in nearly all cases. In the few instances where poor mixing or convergence was noted, we ran additional independent analyses until an adequate sample from the target density could be obtained, or it became clear that the MCMC could not adequately sample from the target distribution. Inferences under each model were based on the combined stationary samples from each of the independent chains, which provided adequate sampling for all parameters according to the effective sample size (ESS) (Drummond et al. 2012).

Finally, we assessed support for the key phylogenetic findings of each published miRNA study using Bayes factors. This entailed a second round of analyses targeting the marginal likelihood density that were identical to our initial analyses under the best fitting clock model (as judged by the Bayes factor model comparisons above), but with the topology constrained to the relevant alternative hypothesis in each case (discussed in more detail below). These analyses allowed us to quantify the extent to which each miRNA dataset can decisively distinguish among alternative phylogenetic hypotheses.

*Patterns and rates of miRNA evolution.*—We used Bayesian model-comparison methods to assess the fit of the miRNA datasets to four (relaxed) clock models, which differ in their ability to accommodate rate variation across lineages. The strict clock makes the most stringent assumption of rate homogeneity, the random-local clock is intermediate, and the uncorrelated (exponential and lognormal) relaxed-clock models are able to capture the most extreme rate fluctuations across branches—rates on adjacent branches are modeled as independent and identically distributed random variables drawn from a common (exponential or lognormal) probability distribution (Drummond et al., 2006). Interestingly, the two uncorrelated relaxed-clock models had the highest marginal likelihood and were therefore the preferred model for every single dataset (Table 2). We were unable to perform a few of these comparisons due to poor mixing of MCMC that prohibited stable estimation of a marginal likelihood for some of the data + model combinations (the uncorrelated lognormal in particular, see Table 2). However, the uncorrelated exponential model was very strongly preferred (2 ln BF > 10) to the Strict model for four datasets, and was strongly preferred (2 ln BF > 6) for the fifth. These results, combined with the large coefficient of variation for rates among branches under the winning model (Table 2), imply substantial heterogeneity in the rate of miRNA evolution across branches in these datasets, conditions in which parsimony inferences are more likely to be inconsistent (*e.g.*, Felsenstein 1978; Huelsenbeck and Hillis 1993; Huelsenbeck 1995). Finally, as in the case of the Dollo parsimony analyses, Bayesian estimates under the stochastic Dollo model indicate substantial rates of miRNA loss in all five miRNA datasets (Table 1).

**Table 2:** Marginal likelihoods of miRNA datasets under four different clock models ranging from strictly clock-like to highly variable evolutionary rates.

| miRNA Study | Marginal Likelihood<sup>a</sup> |  |  |  | CV<sup>b</sup> |
| --- | --- | --- | --- | --- | --- |
|  | Strict | Random local | Uncorrelated exponential | Uncorrelated lognormal |  |
| Amniotes | -128.44<br>(0.06) | -127.93<br>(0.14) | -124.00*<br>(0.02) | -125.22<br>(0.53) | 0.996 |
| Animals | -649.83<br>(0.15) | -639.60<br>(0.36) | -605.37*<br>(0.08) | — | 1.117 |
| Annelids | -454.75<br>(0.16) | — | -433.34*<br>(0.21) | — | 1.105 |
| Bilaterians | -622.88<br>(0.25) | -602.47<br>(0.89) | -583.58*<br>(0.31) | — | 1.107 |
| Vertebrates | -1107.98<br>(0.17) | -1054.95<br>(0.06) | -1034.07*<br>(0.17) | -1037.44<br>(0.38) | 1.043 |

<sup>a</sup>The marginal log probability of miRNA datasets under the stochastic Dollo and (relaxed) clock models estimated using path sampling (stepping stone estimates are nearly identical and are available in the supplementary material). Values are means and standard error of three independent runs. The winning models are denoted with an asterisk. Empty cells denote the model-dataset combinations for which poor MCMC mixing prevented a stable estimate of the marginal likelihood.
<sup>b</sup>Coefficient of Variation in evolutionary rate among branches of the phylogeny for the winning model.

*Evaluating support for key phylogenetic conclusions of published miRNA studies.*—Bayesian analyses of miRNA data offered novel insight into several previously published studies. In three of the five cases, the Bayesian analysis recovers a result that disagrees in important respects from the parsimony result, but agrees with other published studies based on more-traditional phylogenomic analyses of molecular sequence datasets. Parsimony and Bayesian analyses recover congruent conclusions for the two remaining studies, although both of these cases remain problematic due to large uncertainty or sampling error. We briefly discuss key results for each of these analyses below (for additional details, see *Supplemental File 1*).

*Annelid dataset.*—Sperling et al. (2009) sought to evaluate the monophyly of and establish phylogenetic relationships within annelids. Based on the parsimony analysis of the miRNA dataset, they concluded that: 1) annelids are monophyletic (*Nereis*, *Lumbricus*, and *Capitella* form a clade); 2) the sipunculan species, *Phascolosoma*, is the sister group of annelids; and finally, 3) polychaete annelids are not monophyletic (*Nereis* and *Capitella* do not form a clade). Bayesian analysis of the miRNA data under the stochastic Dollo model infers the tree: ((*Nereis*, *Phascolosoma*), (*Lumbricus*, *Capitella*)). Accordingly, these results neither support annelid monophyly nor a sister-group relationship between sipunculans and annelids. Our finding that sipunculids (represented by *Phascolosoma*) are included within annelids—and thus, that annelids are paraphyletic—is consistent with most recent molecular phylogenetic/omic studies (*e.g.*, Colgan et al. 2006; Hausdorf et al. 2007; Rousset et al. 2007; Struck et al. 2007; Dunn et al. 2008; Xin et al. 2009).

We assessed the decisiveness of support for these alternative topological models by performing analyses in which the topology was constrained alternatively to the parsimony estimate (Model *M*_1_, Table 3) and the Bayesian estimate (Model *M*_0_, Table 3) and compared the marginal likelihoods under the two models. A 2 ln BF of *∼* 12 in favor of the Bayesian topology suggests that the data very strongly prefer the Bayesian estimate relative to the parsimony estimate (Kass and Raftery 1995).

**Table 3:** Selection of topology models (tests of phylogenetic hypotheses) for miRNA datasets based on Bayes factor comparisons of estimated marginal likelihoods.

| miRNA study | Topology model<sup>a</sup> | $\ln P(X\vert M_i)^b$ | $2 \ln BF_{ij}^c$ | Description | Resulting from<sup>d</sup> |
| --- | --- | --- | --- | --- | --- |
| Amniotes | M0 | -114.98<br>(0.03) | 17.52 | Lepidosaur hypothesis: turtles + lizards | B & P |
|  | M1 | -123.74<br>(0.14) |  | Archosaur hypothesis: turtles + archosaurs |  |
| Amniotes-corrected<sup>e</sup> | M0 | -126.21<br>(0.22) | 5.17 | Archosaur hypothesis: turtles + archosaurs | B & P |
|  | M1 | -128.80<br>(0.08) |  | Lepidosaur hypothesis: turtles + lizards |  |
| Animals | M0 | -574.67<br>(0.44) | -11.69 | ((Acoel 1, Acoel 2), Xenoturbella), (remaining Bilateria)) | B |
|  | M1 | -568.83<br>(0.17) |  | (Acoel 1 (Acoel 2 (Xenoturbella (remaining Bilateria)))) | P |
| Annelids | M0 | -414.65<br>(0.11) | 11.97 | ((Phascolosoma, Nereis), (Lumbricus, Capitella)) | B |
|  | M1 | -420.64<br>(0.23) |  | (Phascolosoma (Nereis (Lumbricus, Capitella))) | P |
| Bilaterians | M0 | -552.63<br>(0.02) | 100.92 | myzostomids sister to annelids | B |
|  | M1 | -603.09<br>(0.26) |  | myzostomids nested within annelids | P |
| Vertebrates | M0 | -994.81<br>(0.14) | 0.91 | cyclostome hypothesis: lampreys sister to hagfish | B & P |
|  | M1 | -995.26<br>(0.13) |  | jawed vertebrate hypothesis: lampreys sister to jawed vertebrates |  |

<sup>a</sup>Topology models refer to various phylogenetic hypotheses corresponding to the description column; see text for details.
<sup>b</sup>The marginal log probability of miRNA datasets (and standard error) under the stochastic Dollo model and the preferred relaxed-clock model estimated using path sampling as described in the text.
<sup>c</sup>Two times the natural log of the Bayes factor is twice the difference between the natural log marginal likelihoods estimated under the alternative topological models. Our interpretation follows Kass and Raftery (1995).
<sup>d</sup>An indicator for which unconstrained analysis type recovered each topological model. B= Bayesian Stochastic Dollo, P = Parsimony
<sup>e</sup>This is a version of the amniote miRNA dataset from the study of Lyson et al. (2011) that has been corrected for sampling error; see text for details.

*Bilaterian dataset.*—Helm et al. (2012) sought to resolve the phylogenetic affinity of myzostomid worms using an expanded version of the miRNA dataset from the Sperling et al. (2009) study, testing alternative hypotheses that either placed myzostomids within annelids or platyzoans. Their Dollo parsimony analysis of the miRNA data “strongly confirms a phylogenetic position of *Myzostomida*” as “deeply nested within the annelid radiation, as sister to *Capitella*.” By contrast, Bayesian analysis of this miRNA dataset under the stochastic Dollo model implies that myzostomids are the sister group of annelids (with a clade probability of *∼* 0.97–0.99), which agrees with estimates based on recent analyses of phylogenomic data (*e.g.*, Struck et al. 2007).

We assessed the support for these alternative hypotheses by performing analyses in which the topology was constrained to the parsimony estimate (model *M*_1_, Table 3), and compared the marginal likelihood of this model to that from analyses constrained to the Bayesian estimate (model *M*_0_, Table 3). These analyses decisively reject the inclusion of *Myzostoma* within annelids (2 ln BF *∼* 100; Kass and Raftery 1995). It was not possible to perform a clear test of the alternative ‘platyzoan’ hypothesis, as Platyzoa was not inferred to be monophyletic in our unconstrained analyses (for details, see *Supplementary File 1*).

*Animal dataset.*—Philippe et al. (2011) sought to establish the phylogenetic placement of acoels and xenoturbellids within animals using three independent datasets: a large number of mitochondrial genes, a phylogenomic dataset comprising 38, 330 amino-acid positions, and a microRNA dataset. The phylogeny inferred from their Dollo parsimony analysis of the miRNA dataset implied that acoels (*Symsagittifera* and *Hofstenia*) and xenoturbellids (*Xenoturbella*) form a paraphyletic grade near the base of bilaterians: (*Symsagittifera* (*Hofstenia* (*Xenoturbella* (remaining bilaterians)))). The Bayesian analysis of this miRNA dataset under the stochastic Dollo model infers a very different tree in which acoels are monophyletic and sister to xenoturbellids: (((*Symsagittifera*, *Hofstenia*), *Xenoturbella*), remaining bilaterians). We assessed support for these hypotheses by performing additional analyses in which the topology was alternatively constrained to the parsimony estimate (topological model *M*_1_, Table 3) and the Bayesian estimate (topological model *M*_0_, Table 3) and compared the marginal likelihoods. The Bayes factor suggests that the miRNA data favor the parsimony hypothesis (2 ln BF *∼ -*12; Kass and Raftery 1995). Notably however, Philippe et al. (2011) favored a different hypothesis based on their molecular sequence information and expressed caution in interpreting the apparent phylogenetic signal in the miRNA data.

A central result from Philippe et al. (2011) is the close relationship between *Xenoturbella* and (a monophyletic) Acoela (*Symsagittifera*, *Hofstenia*). Although this result strongly conflicts with their parsimony analysis of miRNA data, they prefer it based on their rigorous Bayesian analyses of large-scale molecular datasets. In fact, in discussing the conflicting estimates based on their Bayesian analyses of the phylogenomic data and their parsimony analysis of the miRNA data, Philippe et al. (2011) were skeptical of the miRNA phylogeny, attributing this discrepancy to the effects of pervasive secondary loss of miRNA families in acoels. Interestingly, our Bayesian analysis of the miRNA dataset recovers the same monophyletic Acoela sister to *Xenoturbella*. However, both Bayesian and parsimony analyses of the miRNA data conflict with the preferred tree from Philippe et al. (2011) in other respects, suggesting that secondary loss has strongly obscured any phylogenetic signal in these data.

*Vertebrate dataset.*—Heimberg et al. (2010) sought to resolve the phylogenetic position of lampreys within vertebrates using miRNA data, testing alternative hypotheses that either placed lampreys as sister to hagfish (the ‘cyclostome’ hypothesis) or to jawed vertebrates (the ‘vertebrate’ hypothesis). Analysis of the vertebrate miRNA dataset using Dollo parsimony supported the cyclostome hypothesis: the two lampreys, *Lampetra* and *Petromyzon*, form a clade that is sister to the hagfish species, *Myxine*: ((*Lampetra*, *Petromyzon*), *Myxine*)). Bayesian analysis of the vertebrate miRNA dataset under the stochastic Dollo model also supported the cyclostome hypothesis, albeit weakly (i.e., with a clade probability of *∼* 0.78).

We assessed the support for cyclostome monophyly by performing analyses in which the topology was constrained to the alternative phylogenetic hypothesis in which lampreys are sister to jawed vertebrates (model *M*_1_, Table 3), and compared the marginal likelihoods of the constrained and unconstrained (model *M*_0_, Table 3) analyses. Comparison of the marginal likelihoods under the constrained and unconstrained models suggests that the miRNA data are essentially equivocal regarding the phylogenetic affinity of lampreys (2 ln BF *∼* 1; Kass and Raftery 1995).

*Amniote dataset.*—Lyson et al. (2011) sought to resolve the phylogenetic placement of turtles within amniotes, using a miRNA dataset to test whether turtles were either sister to lizards + tuatara (the ‘lepidosaur’ hypothesis), or to birds + crocodilians (the ‘archosaur’ hypothesis). Analysis of the miRNA dataset using Dollo parsimony supports the lepidosaur hypothesis, and this finding was also strongly supported by Bayesian analysis under the stochastic Dollo model (with a clade probability of *∼* 0.99).

We further assessed support for the lepidosaur hypothesis by performing analyses of the amniote miRNA dataset in which the topology was constrained to the alternative phylogenetic hypothesis in which turtles are sister to archosaurs (model *M*_1_, Table 3), and compared the marginal likelihoods to those from the lepidosaur hypothesis (model *M*_0_, Table 3). In contrast to all other studies, comparison of the marginal likelihoods under the two models suggests that the miRNA data provide strong support for the originally published result (2 ln BF *∼* 17; Kass and Raftery 1995). However, we demonstrate below that this result is an artifact of sampling error in the detection of amniote miRNAs (see: *Sampling error in miRNA detection and its phylogenetic impact*).

*Anomalous results from miRNA analyses.*—Bayesian analysis of published miRNA datasets casts considerable doubt on the key phylogenetic conclusions of those studies. In three of five cases (animals, annelids, and bilaterians), using a model that accounts for the uncertainty in character histories changes the key phylogenetic conclusion, often with strong support. In a fourth case (vertebrates), considering the uncertainty in character history leads to the conclusion that miRNAs are essentially silent on the relationship of interest. In only one case (amniotes) does accounting for uncertainty in character history leave the key conclusion unchanged, although this case reveals a second issue that we explore below. Moreover, our re-analyses of published miRNA datasets also supported some highly unusual phylogenetic results. For example, Bayesian analyses of the amniote miRNA dataset failed to support the (virtually incontrovertible) monophyly of archosaurs, whereas analyses of the animal miRNA dataset supported (the very odd placement of) chordates as the sister to all other bilaterians. We argue below that such remarkable findings likely have a more prosaic explanation.

Shortly after the present manuscript returned from an initial round of peer review, a paper appeared that further discussed the phylogenetic potential of miRNAs and demonstrated phylogenetic inference with miRNAs using the binary stochastic Dollo model (Tarver et al. 2013). This paper assembled a dataset of miRNA presence/absence for 29 metazoan taxa from subsets of the data matrices developed in previous studies (including those that we re-examine here) and analyzed it using the stochastic Dollo. This analysis recovers high posterior probabilities on all nodes except one and is congruent with other phylogenies constructed from more traditional phylogenetic and phylogenomic analyses. Thus, the Tarver et al. (2013) result appears to be in stark contrast with our results. The discrepancy appears to stem from the choice of taxa for inclusion in the Tarver et al. (2013) data matrix. The dataset retains only a subset of the taxa reported in the original studies, while we analyze the original studies’ data matrices in full. Further, the Tarver et al. (2013) matrix is missing all the taxa that we identify as leading to problematic results above. For example, we identify low support and pervasive uncertainty associated with the relationship between the lamprey (*Lampetra* and *Petromyzon*) and the hagfish (*Myxine*)—the central taxa under study in the dataset of Heimberg et al. (2010). Tarver et al. (2013) retain only one lamprey (and no hagfish) from this dataset and thus do not test the support for this clade. Similarly, the acoels (*Symsagittifera*, *Hofstenia*) and *Xenoturbella* are central to the study by Philippe et al. (2011). These taxa disagree strongly with traditionally constructed phylogenies but are not included in Tarver et al. (2013). The two birds (*Gallus* and *Taenopygia*) and lizard from the Lyson et al. (2011) dataset are included in Tarver et al. (2013), but the critical turtle and alligator data are not. Likewise, the key taxon *Myzostomida* from Helm et al. (2012) is not included, nor are *Nereis* and *Phascolosoma* from Sperling et al. (2009). No details outlining the choice of taxa for this matrix are given, so we are unsure why only subsets of previous datasets were included, nor why certain taxa were included versus not. That said, the apparent discrepancy among our results appears to stem from our varying choices of taxa. Because the utility of miRNAs in phylogenetics lies in their purported ability to resolve particularly vexing phylogenetic relationships, our view is that including taxa that allow for tests of such vexing relationships is a critical part of studying these marker’s phylogenetic utility.

## Sampling error in miRNA detection and its phylogenetic impact

Sampling error can to lead to the (apparent) absence of miRNAs in phylogenetic datasets. This is of particular concern because most miRNA phylogenetic studies use a mixture of approaches to identify miRNAs in different lineages (namely, using a combination of bioinformatic scans of complete genomes and/or *de novo* sequencing of small-RNA libraries). If these approaches vary in their detection probabilities, then miRNAs are more likely to be discovered in some lineages than in others. As more and more data are collected under this biased detection scheme, certain lineages are likely to accumulate true presences while the remaining lineages will accumulate apparent absences. Since the presence and absence of miRNAs are the direct source of phylogenetic information, this sampling artifact may lead to biased estimates of topology.

Here we demonstrate sources of sampling error in the detection of miRNA families, first focusing on the analysis of turtle relationships within amniotes as a detailed case study, and then assessing the generality of this sampling error by means of a more general empirical survey.

*Sampling bias in the detection of amniote miRNAs.*—Lyson et al. (2011) employed a mixture of miRNA detection methods in an attempt to resolve the phylogenetic position of turtles within amniotes. Specifically, their study searched for miRNAs using: 1) similarity searches against whole-genome assemblies for two birds—chicken (*Gallus*), zebra finch (*Taeniopygia*)—and four outgroup taxa; 2) a combination of similarity searches against the genome assembly for the lizard (*Anolis*) and *de novo* sequencing of an *Anolis* RNA library; and 3) *de novo* sequencing of RNA libraries for a turtle species—the painted turtle (*Chrysemys*)—and the American alligator (*Alligator*). At the time of their study, full genome assemblies for the painted turtle and alligator were not available. The authors identified 19 miRNA families unique to birds, one miRNA family unique to archosaurs (birds and crocodilians), but no miRNA families shared between archosaurs and turtles. Furthermore, the study identified four miRNA families that are shared between the anole and turtle. Taken at face value, these data appear to unequivocally support a turtle + lizard relationship, to the exclusion of archosaurs.

Draft genome assemblies for both the painted turtle and American alligator are now available (St John et al. 2012; Shaffer et al. 2013), which provide an independent check of the miRNAs detected—and the phylogenetic conclusions reached—in the Lyson et al. (2011) study. We sought to confirm that each of the miRNA families that were identified by Lyson et al. (2011) as unique to birds (*N* = 19) were in fact absent from the turtle and alligator genomes, and that the single archosaur-specific miRNA was absent from the turtle genome. We also assessed whether each of the miRNA families that were identified as being shared exclusively by turtles and lizards were in fact present in the turtle genome and absent from the alligator genome.

We downloaded both the longer stem-loop sequence (60–80 bp) and the shorter mature sequence (22 bp) for each relevant miRNA from miRBase (Kozomara and Griffiths-Jones 2011) for each appropriate reference taxon (*Gallus* for the 19 bird-specific and the single archosaur-specific miRNA families; *Anolis* for the four miRNA families uniquely shared by turtle + lizard). We constructed local BLAST databases from the turtle and alligator genome assemblies (*v*3.0.3 and 0.1*d*27, respectively) and searched against them with each of the relevant miRNA stem-loop sequences using BLASTN (*v*2.2.25, minimum word size = 11, e-value cutoff = 10–2; Zhang et al. 2000). We then predicted secondary structure for any putative miRNAs that we identified using mFold (Zuker 2003).

We scored a miRNA family as being present in the turtle and/or alligator genome if it met three criteria: 1) We observed a highly significant hit (*i.e.*, with a minimum e-value of 10*^-^*^20^) for the reference stem-loop sequence against the relevant genome assembly; 2) The matching sequence in the genome contained a nearly perfect match to the mature *∼*22 bp miRNA sequence (*i.e.*, containing no more than one substitution in the mature miRNA sequence); 3) The matching sequence in the turtle or alligator genome folded into the expected hairpin secondary structure and this structure was similar to the predicted secondary structure published for the reference sequence.

Our search confirmed that the single archosaur-specific miRNA (miRNA 1791 in Lyson et al. 2011) was present in the alligator genome, as expected. However, we discovered that this miRNA is also present in the turtle genome (for sequences and predicted secondary structure, see *Supplementary File 2*). Furthermore, we discovered three additional miRNA families present in both the alligator and turtle genomes that were reported by Lyson et al. (2011) as being unique to birds (miRNA families 1641, 1743, and 2964). All four families exhibited very high sequence similarity with the known miRNA from the reference taxon, highly conserved stem-loop structures with similar free energies to that predicted from the reference taxon, and mature sequences that were identical (two families) or nearly identical (two families) to the reference (see *Supplementary File 2* for sequence alignments and predicted structures). This sampling error may be inherent to miRNA-detection approaches that rely on RNA sequencing. For example, Sperling et al. (2009) observed a similar pattern in the polychaete worm, *Capitella*. They discovered five additional miRNAs from the genome of this organism that were not detected in the sequences derived from an RNA library. MicroRNAs are frequently expressed only in certain tissues, at certain stages of development, or expressed at low levels (Sperling et al. 2009; Landgraf et al. 2007; Powder et al. 2012; Darnell et al. 2006; Wienholds et al. 2005). In these cases, it is likely that miRNAs actually present in the genome will be missed because they are not being transcribed (or only being transcribed at low levels) in the tissue that was used to make the RNA library.

Finally, we sought to confirm that the four miRNA families identified by Lyson et al. (2011) as uniting a lizard + turtle clade were, in fact, present in the turtle genome and absent in the alligator genome (miRNA families 5390, 5391, 5392, and 5393). Our search confirmed that all four miRNA families were absent from the alligator genome, as expected. However, we were only able to find one of the four reported miRNA families (miRNA 5391) in the turtle genome. We found no significant BLAST hits to any of the other three expected miRNAs, even under relaxed search settings (word size = 4, e-value cutoff = 10). We then assessed whether we could identify these miRNAs in the *Anolis* genome and found all four families, as expected. At present, the cause of this discrepancy is unclear. Our failure to detect these sequences could be a false negative, indicating that the turtle genome assembly is incomplete and missing these three sequences. Alternatively, their previous detection could be a false positive in the Lyson et al. (2011) study, stemming from contamination between the *Anolis* and *Chrysemys* sequencing libraries or from another source of error. The turtle genome assembly has 18x coverage and is estimated to be 93% complete, which suggests that the former explanation is unlikely (Shaffer et al. 2013). Nevertheless, we can not formally distinguish between these possibilities at present.

We then revised the Lyson et al. (2011) data matrix to correct this sampling error and subjected the revised matrix to Bayesian phylogenetic analysis under the stochastic Dollo model (analyses performed as detailed above). Rather than supporting a strong relationship between lizards and turtles, the corrected miRNA dataset supports a relationship between turtles and archosaurs, albeit weakly (i.e., with a clade probability of *∼* 0.54). This result is consistent with several recently published studies that examine the phylogenetic placement of turtles using large DNA sequence datasets (Shaffer et al. 2013; Crawford et al. 2012; Shen et al. 2011; Chiari et al. 2012).

We assessed support for the ‘archosaur’ hypothesis by performing analyses of the corrected amniote miRNA dataset in which the topology was constrained to the alternative ‘archosaur’ and ‘lepidosaur’ hypotheses (models *M*_0_ and *M*_1_ in Table 3, respectively). Comparison of the marginal likelihoods under the alternative models indicate that the miRNA data provide positive evidence in favor of the archosaur hypothesis (2 ln BF *∼* 5). This analysis illustrates that miRNA detection is prone to strong sampling error, to a degree that can fundamentally alter the conclusions of phylogenetic inferences based on these data.

*General survey of sampling bias in miRNA detection.*—Our ability to provide a detailed description of the miRNA detection bias in the amniote study largely rests on the serendipitous availability of two new genome assemblies. Accordingly, it is not possible to perform a comparably detailed analysis of the potential sampling errors in the other four published miRNA phylogenetic studies. However, we can make a more general comparison of alternative miRNA detection strategies. To do so, we compiled information from the literature of cases in which the total miRNA complement of various organisms had been estimated both by means of *de novo* sequencing of small-RNA libraries and also by means of bioinformatic searches of DNA sequence resources. If no sampling bias exists, of course, (virtually) identical sets of miRNA families should be identified using alternative strategies. In stark contrast to this expectation, however, we see a high degree of variation in the miRNA complement identified under the two strategies (Table 4). Although this comparison does not directly replicate the alternative methods employed in published phylogenetic studies, it clearly indicates the prevalence of variation in total miRNA complement detection and, as we have shown, this type of sampling error has the potential to impact estimates of phylogeny.

**Table 4:** Comparison of empirical and computationally derived estimates of miRNA complements for selected taxa.

| Species | Number of<br>miRNA orthologs<br>obtained<br>empirically | Number of miRNA orthologs accessioned in: |  |
| --- | --- | --- | --- |
|  |  | miROrtho<sup>a</sup> | miRBase<sup>b</sup> |
| *Apis mellifera* | 267<sup>c</sup> | 52 | 222 |
| *Tribolium castaneum* | 203<sup>d</sup> | 35 | 430 |
| *Drosophila melanogaster* | 148<sup>e</sup> | 147 | 426 |
| *Caenorhabditis elegans* | 112<sup>f</sup> | 130 | 368 |
| *Schmidtea mediterranea* | 122<sup>g</sup>; 66<sup>h</sup> | 38 | 257 |
| *Schistosoma japonicum* | 227<sup>i</sup> | - | 78 |
| *Schistosoma mansoni* | 211<sup>j</sup> | - | 29 |
| *Petromyzon marinus* | 267<sup>k</sup> | 40 | 302 |
| *Branchiostoma floridae* | 152<sup>k</sup>; 32<sup>l</sup> | - | 187 |
| *Saccoglossus kowalevskii* | 90<sup>k</sup> | - | 115 |
| *Strongylocentrotus purpuratus* | 58<sup>k</sup> | 12 | 70 |
| *Danio rerio* | 198<sup>m</sup> | 113 | 255 |
| *Oryzias latipes* | 599<sup>n</sup> | - | 146 |
| *Nematostella vectensis* | 40<sup>o</sup> | - | 78 |

<sup>a</sup>Gerlach et al. (2009)
<sup>b</sup>Kozomara and Griffiths-Jones (2011)
<sup>c</sup>Chen et al. (2010)
<sup>d</sup>Marco et al. (2010)
<sup>e</sup>Ruby et al. (2007)
<sup>f</sup>de Wit et al. (2009)
<sup>g</sup>Friedländer et al. (2009)
<sup>h</sup>Lu et al. (2009)
<sup>i</sup>Xue et al. (2008)
<sup>j</sup>Simões et al. (2011)
<sup>k</sup>Campo-Paysaa et al. (2011)
<sup>l</sup>Dai et al. (2009)
<sup>m</sup>Soares et al. (2009)
<sup>n</sup>Li et al. (2010)

## Conclusions

The current wealth of molecular data will continue to resolve relationships in the tree of life, but not all nodes will acquiesce with equal effort. Predictably, the variously recalcitrant, enigmatic, inscrutable and impenetrable relationships will continue to increase in prevalence. Ultimately, resolution of these problematic cases may require the discovery of new and improved phylogenetic data (and/or the elaboration and careful application of more realistic models that better describe important aspects of the processes that give rise to conventional genomic data). Accordingly, it is predictable that the addition of a putative silver bullet—such as miRNA presence/absence data—to our phylogenetic arsenal will be greeted with enthusiasm. We would argue, however, that this enthusiasm should be tempered with careful consideration of how to appropriately accommodate the correspondingly novel processes by which these new data evolved and/or new procedures by which they are collected.

We have demonstrated that the evolution of miRNA families is apparently complex. Contrary to repeated claims, secondary loss of miRNA appears to be quite prevalent, and miRNA evolution typically exhibits substantial variation in rate across branches through time. Consequently, the complex character histories associated with miRNA evolution suggest that parsimony—which effectively places all of the probability on the character history with the minimal change—is not a defensible method with which to infer phylogeny from these new data. We have demonstrated that, in principle, it is both possible and preferable to estimate phylogeny from miRNA data within a Bayesian statistical framework using stochastic evolutionary models. Adopting a statistical approach for estimating phylogeny from miRNA (or other) data confers many benefits: this approach allows us to choose objectively among models, to perform formal tests of competing hypotheses, promotes a richer study of the evolutionary process, and enables us to gauge and accommodate uncertainty in our estimates. We have established the importance of adopting a more appropriate statistical approach: Bayesian analyses of published miRNA datasets qualitatively altered key phylogenetic conclusions and/or revealed considerable phylogenetic uncertainty in these estimates in four of the five cases that we examined.

Finally, we have demonstrated that the detection of miRNA families is prone to error—especially when using a mixture of detection methods—and this sampling error can substantially bias estimates of phylogeny. Accordingly, it is critical that we either extend existing stochastic models to accommodate this ascertainment bias, or take precautionary measures to minimize it. For example, models used to analyze both SNP data in population genetics (Clark et al. 2005) and discrete-morphological data in phylogenetics (Ronquist et al. 2012) explicitly model the associated ascertainment strategies in order to reduce the associated biases. The stochastic Dollo model might be similarly extended to accommodate the documented miRNA ascertainment bias. However, the complexity of the mixed genomic/RNA-library detection strategy would make such an extension challenging, although the intense focus on miRNA detection methods (*e.g.*, Pritchard et al. 2012) gives reason for optimism that these extensions may be possible. Alternatively, studies seeking to estimate phylogeny from miRNA presence/absence data should strictly employ identical, genome-based detection methods in all lineages. This may not always eliminate sampling error, but it should reduce bias arising from differential detection probabilities of the various miRNA discovery methods.

Although our appraisal of miRNA as a novel source of phylogenetic information is admittedly critical, we clearly recognize the potential of these data to inform phylogeny: inferences based on miRNA data often correspond broadly to those based on more conventional gene/omic data. We take issue, however, with the recent promotion of miRNA data as a phylogenetic panacea. New data are attended by new issues that need to be carefully resolved in order to realize their full potential.

## Acknowledgments

We thank Artyom Kopp and the members of a phylogenetic reading group at UC Davis for helpful discussion and advice during the development of this project. We also thank the Turtle Genome Sequencing Consortium and the International Crocodilian Genomes Working Group for providing pre-publication access to the genome assemblies used in this study. Support for this work was provided in part by National Science Foundation grants DEB-0842181 and DEB-0919529 to BRM.

### Supplementary Material

Supplementary material may be found in the Dryad data repository at http://datadryad.org.

Supplementary File 1—Workbook with details of the Bayesian phylogenetic analyses of the miRNA datasets.

Supplementary File 2—Sequence alignment and predicted secondary structure for four microRNA families that were detected in the *Alligator* and *Chrysemys* genomes via BLAST similarity searches. The mature miRNA sequence from miRBase is underlined in the sequence and secondary structure of the reference species (*Gallus*). Substitutions relative to the reference sequence are highlighted in red. miRNA 1743 sits at the end of a contig in the *Chrysemys* genome assembly and is truncated by 10 bases on the 5’ end as a result. We represent these as ambiguous bases and make no attempt to predict secondary structure in this region.

## Footnotes

1 Several additional studies discuss the phylogenetic implications of miRNA data, but do not subject these data to a formal phylogenetic analysis. Typically in these studies, the phylogeny is first estimated from some other source of data, and then the correspondence of the inferred tree to select miRNA families is discussed (e.g., Rota-Stabelli et al. 2011; Sempere et al. 2007; Wheeler et al. 2009; Campbell et al. 2011; Sperling et al. 2011).

## References

Alekseyenko, A. V., C. J. Lee, and M. A. Suchard. 2008. Wagner and Dollo: A stochastic duet by composing two parsimonious solos. Systematic Biology 57:772–784.

Alvarez-Garcia, I. and E. A. Miska. 2005. MicroRNA functions in animal development and human disease. Development (Cambridge, England) 132:4653–62.

Baele, G., P. Lemey, T. Bedford, A. Rambaut, M. Suchard, and A. V. Alekseyenko. 2012. Improving the accuracy of demographic and molecular clock model comparison while accommodating phylogenetic uncertainty. Molecular Biology and Evolution 29:2157–67.

Berezikov, E. 2011. Evolution of microRNA diversity and regulation in animals. Nature Reviews Genetics 12:846–60.

Boore, J. L. 2006. The use of genome-level characters for phylogenetic reconstruction. Trends in Ecology & Evolution 21:439–446.

Boore, J. L. and S. I. Fuerstenberg. 2008. Beyond linear sequence comparisons: The use of genome-level characters for phylogenetic reconstruction. Philosophical Transactions of the Royal Society of London. Series B, Biological Sciences 363:1445–1451.

Bremer, K. 1988. The limits of amino acid sequence data in angiosperm phylogenetic reconstruction. Evolution 42:795–803.

Campbell, L. I., O. Rota-Stabelli, G. D. Edgecombe, T. Marchioro, S. J. Longhorn, M. J. Telford, H. Philippe, L. Rebecchi, K. J. Peterson, and D. Pisani. 2011. MicroRNAs and phylogenomics resolve the relationships of Tardigrada and suggest that velvet worms are the sister group of arthropoda. Proceedings of the National Academy of Science, USA.

Campo-Paysaa, F., M. Sémon, R. A. Cameron, K. J. Peterson, and M. Schubert. 2011. MicroRNA complements in deuterostomes: Origin and evolution of microRNAs. Evolution & Development 13:15–27.

Chen, X., X. Yu, Y. Cai, H. Zheng, D. Yu, G. Liu, Q. Zhou, S. Hu, and F. Hu. 2010. Next-generation small RNA sequencing for microRNAs profiling in the honey bee *Apis mellifera*. Insect molecular biology 19:799–805.

Chiari, Y., V. Cahais, N. Galtier, and F. Delsuc. 2012. Phylogenomic analyses support the position of turtles as the sister group of birds and crocodiles (Archosauria). BMC Biology 10:65.

Clark, A., M. J. Hubisz, C. D. Bustamante, S. H. Williamson, and R. Nielsen. 2005. Ascertainment bias in studies of human genome-wide polymorphism. Genome Research 15:1496–1502.

Colgan, D. J., P. A. Hutchings, and M. Braune. 2006. A multi-gene framework for polychaete phylogenetic studies. Organismal Diversity & Evolution 6:220–235.

Crawford, N. G., B. C. Faircloth, J. E. McCormack, R. T. Brumfield, K. Winker, and T. C. Glenn. 2012. More than 1000 ultraconserved elements provide evidence that turtles are the sister group of archosaurs. Biology Letters.

Dai, Z., Z. Chen, H. Ye, L. Zhou, L. Cao, Y. Wang, S. Peng, and L. Chen. 2009. Characterization of microRNAs in cephalochordates reveals a correlation between microRNA repertoire homology and morphological similarity in chordate evolution. Evolution & development 11:41–49.

Darnell, D. K., S. Kaur, S. Stanislaw, J. K. Konieczka, T. A. Yatskievych, and P. B. Antin. 2006. MicroRNA expression during chick embryo development. Developmental Dynamics 235:3156–3165.

de Wit, E., S. E. V. Linsen, E. Cuppen, and E. Berezikov. 2009. Repertoire and evolution of miRNA genes in four divergent nematode species. Genome research 19:2064–74.

Dolgin, E. 2012. Phylogeny: Rewriting evolution. Nature 486:460–462.

Drummond, A. J., S. Y. Ho, M. J. Phillips, and A. Rambaut. 2006. Relaxed phylogenetics and dating with confidence. PLoS Biology 4:e88.

Drummond, A. J. and M. A. Suchard. 2010. Bayesian random local clocks, or one rate to rule them all. BMC Biology 8:114.

Drummond, A. J., M. A. Suchard, D. Xie, and A. Rambaut. 2012. Bayesian phylogenetics with BEAUti and the BEAST 1.7. Molecular Biology and Evolution 29:1969–1973.

Dunn, C. W., A. Hejnol, D. Q. Matus, K. Pang, W. E. Browne, S. A. Smith, E. Seaver, G. W. Rouse, M. Obst, G. D. Edgecombe, M. V. Sorensen, S. H. D. Haddock, A. Schmidt-Rhaesa, A. Okusu, R. M. Kristensen, W. C. Wheeler, M. Q. Martindale, and G. Giribet. 2008. Broad phylogenomic sampling improves resolution of the animal tree of life. Nature 452:745–749.

Felsenstein, J. 1978. Cases in which parsimony or compatibility methods will be positively misleading. Systematic Zoology 27:401–410.

Felsenstein, J. 1993. PHYLIP: (Phylogeny Inference Package). Distributed by author.

Friedländer, M. R., C. Adamidi, T. Han, S. Lebedeva, T. A. Isenbarger, M. Hirst, M. Marra, C. Nusbaum, W. L. Lee, J. C. Jenkin, A. Sánchez Alvarado, J. K. Kim, and N. Rajewsky. 2009. High-resolution profiling and discovery of planarian small RNAs. Proceedings of the National Academy of Sciences of the United States of America 106:11546–51.

Gerlach, D., E. V. Kriventseva, N. Rahman, C. E. Vejnar, and E. M. Zdobnov. 2009. miROrtho: computational survey of microRNA genes. Nucleic acids research 37:D111–7.

Grimson, A., M. Srivastava, B. Fahey, B. J. Woodcroft, H. R. Chiang, N. King, B. M. Degnan, D. S. Rokhsar, and D. P. Bartel. 2008. Early origins and evolution of microRNAs and Piwi-interacting RNAs in animals. Nature 455:1193–7.

Guerra-Assunçáo, J. A. and A. J. Enright. 2012. Large-scale analysis of microRNA evolution. BMC genomics 13:218.

Hausdorf, B., M. Helmkampf, A. Meyer, A. Witek, H. Herlyn, I. Bruchhaus, T. Hankeln, and T. S. and B. Lieb. 2007. Spiralian phylogenomics supports the resurrection of Bryozoa comprising Ectoprocta and Entoprocta. Molecular Biology and Evolution 24:2723–2729.

Heimberg, A. M., R. Cowper-Sal-lari, M. Sémon, P. C. J. Donoghue, and K. J. Peterson. 2010. MicroRNAs reveal the interrelationships of hagfish, lampreys, and gnathostomes and the nature of the ancestral vertebrate. Proceedings of the National Academy of Science, USA 107:19379–19383.

Heimberg, A. M., L. F. Sempere, V. N. Moy, P. C. J. Donoghue, and K. J. Peterson. 2008. MicroRNAs and the advent of vertebrate morphological complexity. Proceedings of the National Academy of Sciences of the United States of America 105:2946–50.

Helm, C., S. H. Bernhart, C. H. zu Siederdissen, B. Nickel, and C. Bleidorn. 2012. Deep sequencing of small RNAs confirms an annelid affinity of Myzostomida. Molecular Phylogenetics and Evolution 64:198–203.

Hillis, D. 1999. SINEs of the perfect character. Proceedings of the National Academy of Science, USA 96:9979–9981.

Huelsenbeck, J. P. 1995. Performance of phylogenetic methods in simulation. Systematic Biology 44:17–48.

Huelsenbeck, J. P. and D. M. Hillis. 1993. Success of phylogenetic methods in the four-taxon case. Systematic Biology 42:247–264.

Kass, R. E. and A. E. Raftery. 1995. Bayes factors. Journal of the American Statistical Association 90:773–795.

Kluge, A. G. and J. S. Farris. 1969. Quantitative phyletics and the evolution of anurans. Systematic Zoology 18:1–32.

Kozomara, A. and S. Griffiths-Jones. 2011. miRBase: integrating microRNA annotation and deep-sequencing data. Nucleic acids research 39:D152–7.

Krol, J., I. Loedige, and W. Filipowicz. 2010. The widespread regulation of microRNA biogenesis, function and decay. Nature Reviews Genetics 11:597–610.

Landgraf, P., M. Rusu, R. Sheridan, A. Sewer, N. Iovino, A. Aravin, S. Pfeffer, A. Rice, A. O. Kamphorst, M. Landthaler, C. Lin, N. D. Socci, L. Hermida, V. Fulci, S. Chiaretti, R. FoÃă, J. Schliwka, U. Fuchs, A. Novosel, R.-U. MÃĳller, B. Schermer, U. Bissels, J. Inman, Q. Phan, M. Chien, D. B. Weir, R. Choksi, G. D. Vita, D. Frezzetti, H.-I. Trompeter, V. Hornung, G. Teng, G. Hartmann, M. Palkovits, R. D. Lauro, P. Wernet, G. Macino, C. E. Rogler, J. W. Nagle, J. Ju, F. N. Papavasiliou, T. Benzing, P. Lichter, W. Tam, M. J. Brownstein, A. Bosio, A. Borkhardt, J. J. Russo, C. Sander, M. Zavolan, and T. Tuschl. 2007. A mammalian microRNA expression atlas based on small RNA library sequencing. Cell 129:1401–1414.

LeQuesne, W. J. 1974. The uniquely evolved character concept and its cladistic application. Systematic Zoology 23:513–517.

Li, S.-C., W.-C. Chan, M.-R. Ho, K.-W. Tsai, L.-Y. Hu, C.-H. Lai, C.-N. Hsu, P.-P. Hwang, and W.-c. Lin. 2010. Discovery and characterization of medaka miRNA genes by next generation sequencing platform. BMC genomics 11 Suppl 4:S8.

Lu, J., G. Getz, E. A. Miska, E. Alvarez-Saavedra, J. Lamb, D. Peck, A. Sweet-Cordero, B. L. Ebert, R. H. Mak, A. A. Ferrando, J. R. Downing, T. Jacks, H. R. Horvitz, and T. R. Golub. 2005. MicroRNA expression profiles classify human cancers. Nature 435:834–8.

Lu, Y.-C., M. Smielewska, D. Palakodeti, M. T. Lovci, S. Aigner, G. W. Yeo, and B. R. Graveley. 2009. Deep sequencing identifies new and regulated microRNAs in *Schmidtea mediterranea*. RNA (New York, N.Y.) 15:1483–91.

Lyson, T. R., E. A. Sperling, A. M. Heimberg, J. A. Gauthier, B. L. King, and K. J. Peterson. 2011. MicroRNAs support a turtle + lizard clade. Biology Letters.

Lyu, Y., Y. Shen, H. Li, Y. Chen, L. Guo, Y. Zhao, E. Hungate, S. Shi, C.-I. Wu, and T. Tang. 2014. New microRNAs in Drosophila–birth, death and cycles of adaptive evolution. PLoS genetics 10:e1004096.

Marco, A., J. H. L. Hui, M. Ronshaugen, and S. Griffiths-Jones. 2010. Functional shifts in insect microRNA evolution. Genome biology and evolution 2:686–96.

Meunier, J., F. Lemoine, M. Soumillon, A. Liechti, M. Weier, K. Guschanski, H. Hu, P. Khaitovich, and H. Kaessmann. 2013. Birth and expression evolution of mammalian microRNA genes. Genome research 23:34–45.

Nicholls, G. and R. Gray. 2008. Dated ancestral trees from binary trait data and their application to the diversification of languages. Journal of the Royal Statistical Society, B 70:545–566.

Nozawaet, M., S. Miura, and M. Nei. 2010. Origins and evolution of microRNA genes in drosophila species. Genome Biology and Evolution 2:180–189.

Nylander, J., J. C. Wilgenbusch, D. L. Warren, and D. L. Swofford. 2008. AWTY (are we there yet?): a system for graphical exploration of MCMC convergence in Bayesian phylogenetics. Bioinformatics 24:581.

Peterson, K. J., M. R. Dietrich, and M. A. McPeek. 2009. MicroRNAs and metazoan macroevolution: insights into canalization, complexity, and the Cambrian explosion. BioEssays 31:736–47.

Philippe, H., H. Brinkmann, R. R. Copley, L. L. Moroz, H. Nakano, A. J. Poustka, A. Wallberg, K. J. Peterson, and M. J. Telford. 2011. Acoelomorph flatworms are deuterostomes related to *Xenoturbella*. Nature 470:255–258.

Powder, K. E., Y.-C. Ku, S. A. Brugmann, R. A. Veile, N. A. Renaud, J. A. Helms, and M. Lovett. 2012. A cross-species analysis of microRNAs in the developing avian face. PLoS ONE 7:e35111.

Pritchard, C. C., H. H. Cheng, and M. Tewari. 2012. MicroRNA profiling: Approaches and considerations. Nature Reviews Genetics 13:358–369.

Rambaut, A. and A. J. Drummond. 2007. Tracer v1.4. http://beast.bio.ed.ac.uk/Tracer.

Rokas, A. and P. W. H. Holland. 2000. Rare genomic changes as a tool for phylogenetics. Trends in Ecology & Evolution 15:454–459.

Ronquist, F., M. Teslenko, P. van der Mark, D. L. Ayres, A. Darling, S. Höhna, B. Larget, L. Liu, M. A. Suchard, and J. P. Huelsenbeck. 2012. MrBayes 3.2: Efficient Bayesian phylogenetic inference and model choice across a large model space. Systematic Biology 61:539–542.

Rota-Stabelli, O., L. Campbell, H. Brinkmann, G. D. Edgecombe, S. J. Longhorn, K. J. Peterson, D. Pisani, H. Philippe, and M. J. Telford. 2011. A congruent solution to arthropod phylogeny: Phylogenomics, microRNAs and morphology support monophyletic Mandibulata. Proceedings of the Royal Society: Biological Sciences 278:298–306.

Rousset, V., F. Pleijel, G. W. Rouse, C. Erséus, and M. E. Siddall. 2007. A molecular phylogeny of annelids. Cladistics 23:41–63.

Ruby, J. G., A. Stark, W. K. Johnston, M. Kellis, D. P. Bartel, and E. C. Lai. 2007. Evolution, biogenesis, expression, and target predictions of a substantially expanded set of *Drosophila* microRNAs. Genome research 17:1850–64.

Sanderson, M. J. 2008. Phylogenetic signal in the eukaryotic tree of life. Science 321:121–3.

Sempere, L. F., P. Martinez, C. Cole, J. Baguñá, and K. J. Peterson. 2007. Phylogenetic distribution of microRNAs supports the basal position of acoel flatworms and the polyphyly of Platyhelminthes. Evolution & Development 9:409–415.

Shaffer, H. B., P. Minx, D. E. Warren, A. M. Shedlock, R. C. Thomson, N. Valenzuela, J. Abramyan, C. T. Amemiya, D. Badenhorst, K. K. Biggar, G. M. Borchert, C. W. Botka, R. M. Bowden, E. L. Braun, A. M. Bronikowski, B. G. Bruneau, L. T. Buck, B. Capel, T. A. Castoe, M. Czerwinski, K. D. Delehaunty, S. V. Edwards, C. C. Fronick, M. K. Fujita, L. Fulton, T. A. Graves, R. E. Green, W. Haerty, R. Hariharan, O. Hernandez, L. W. Hillier, A. K. Holloway, D. Janes, F. J. Janzen, C. Kandoth, L. Kong, A. J. de Koning, Y. Li, R. Literman, S. E. McGaugh, L. Mork, M. O’Laughlin, R. T. Paitz, D. D. Pollock, C. P. Ponting, S. Radhakrishnan, B. J. Raney, J. M. Richman, J. St John, T. Schwartz, A. Sethuraman, P. Q. Spinks, K. B. Storey, N. Thane, T. Vinar, L. M. Zimmerman, W. C. Warren, E. R. Mardis, and R. K. Wilson. 2013. The western painted turtle genome, a model for the evolution of extreme physiological adaptations in a slowly evolving lineage. Genome biology 14:R28.

Shen, X.-X., D. Liang, J.-Z. Wen, and P. Zhang. 2011. Multiple genome alignments facilitate development of NPCL markers: A case study of tetrapod phylogeny focusing on the position of turtles. Molecular Biology and Evolution 28:3237–3252.

Simões, M. C., J. Lee, A. Djikeng, G. C. Cerqueira, A. Zerlotini, R. A. da Silva-Pereira, A. R. Dalby, P. LoVerde, N. M. El-Sayed, and G. Oliveira. 2011. Identification of *Schistosoma mansoni* microRNAs. BMC genomics 12:47.

Soares, A. R., P. M. Pereira, B. Santos, C. a. Egas, A. C. Gomes, J. Arrais, J. L. Oliveira, G. R. Moura, and M. A. S. Santos. 2009. Parallel DNA pyrosequencing unveils new zebrafish microRNAs. BMC genomics 10:195.

Sperling, E. A. and K. J. Peterson. 2009. Exchangeability and related topics. Pages 157–170 *in* Animal Evolution–Genomes, Trees and Fossils Oxford Univ Press.

Sperling, E. A., D. Pisani, and K. J. Peterson. 2011. Molecular paleobiological insights into the origin of the Brachiopoda. Evolution and Development 13:290–303.

Sperling, E. A., J. Vinther, B. M. Wheeler, M. Sémon, D. E. G. Briggs, and K. J. Peterson. 2009. MicroRNAs resolve an apparent conflict between annelid systematics and their fossil record. Proceedings of the Royal Society: Biological Sciences 276:4315–4322.

St John, J., E. Braun, S. Isberg, L. Miles, A. Chong, J. Gongora, P. Dalzell, C. Moran, B. Bed’Hom, A. Abzhanov, S. Burgess, A. Cooksey, T. Castoe, N. Crawford, L. Densmore, J. Drew, S. Edwards, B. Faircloth, M. Fujita, M. Greenwold, F. Hoffmann, J. Howard, T. Iguchi, D. Janes, S. Khan, S. Kohno, A. J. de Koning, S. Lance, F. McCarthy, and J. McCormack. 2012. Sequencing three crocodilian genomes to illuminate the evolution of archosaurs and amniotes. Genome Biology 13:415.

Struck, T. H., N. Schult, T. Kusen, E. Hickman, C. Bleidorn, D. McHugh, and K. M. Halanych. 2007. Annelid phylogeny and the status of Sipuncula and Echiura. BMC Evolution Biology 7:57.

Swofford, D. L. 1998. PAUP*: Phylogenetic analysis using parsimony and other methods. Sinauer Associates, Inc., Sunderland, Massachusetts.

Tarver, J. E., E. A. Sperling, A. Nailor, A. M. Heimberg, J. M. Robinson, B. L. King, D. Pisani, P. C. J. Donoghue, and K. J. Peterson. 2013. miRNAs: small genes with big potential in metazoan phylogenetics. Molecular Biology and Evolution 30:2369–82.

Thomson, R. C. and H. B. Shaffer. 2010. Rapid progress on the vertebrate tree of life. BMC Biology 8:19.

Wheeler, B. M., A. M. Heimberg, V. N. Moy, E. A. Sperling, T. W. Holstein, S. Heber, and K. J. Peterson. 2009. The deep evolution of metazoan microRNAs. Evolution & Development 11:50–68.

Wienholds, E., W. P. Kloosterman, E. Miska, E. Alvarez-Saavedra, E. Berezikov, E. de Bruijn, H. R. Horvitz, S. Kauppinen, and R. H. A. Plasterk. 2005. MicroRNA expression in zebrafish embryonic development. Science (New York, N.Y.) 309:310–1.

Xin, S., J. R. X. Ma, and F. Zhao. 2009. A close phylogenetic relationship between Sipuncula and Annelida evidenced from the complete mitochondrial genome sequence of *Phascolosoma esculenta*. BMC Genommics 10:36.

Xue, X., J. Sun, Q. Zhang, Z. Wang, Y. Huang, and W. Pan. 2008. Identification and characterization of novel microRNAs from *Schistosoma japonicum*. PloS one 3:e4034.

Zhang, Z., S. Schwartz, L. Wagner, and W. Miller. 2000. A greedy algorithm for aligning DNA sequences. Journal of Computational Biology 7:203–214.

Zuker, M. 2003. Mfold web server for nucleic acid folding and hybridization prediction. Nucleic Acids Research 31:3406–3415.

